# Micromolar concentrations of metabolites enable coexistence of bacterial species in chemostats

**DOI:** 10.64898/2026.04.23.720424

**Authors:** Eric Ulrich, Sara Mitri

## Abstract

Competition for a single limiting resource is expected to lead to competitive exclusion, yet diverse microbial communities persist even in nutrient-poor environments. Cross-feeding of essential metabolites is one mechanism that can promote coexistence between species, but its contribution is difficult to pinpoint experimentally. Here, we studied a prototroph-auxotroph pair growing on a single carbon source in chemostats. In minimal medium, the prototroph *Comamonas testosteroni* (Ct) supplied thiamine to the thiamine-auxotroph *Ochrobactrum anthropi* (Oa), allowing stable coexistence in agreement with consumer-resource theory. Contrary to our expectation that supplying thiamine would remove the dependency and lead to exclusion of Ct, coexistence persisted even when thiamine was supplemented. Our theoretical anlaysis showed that coexistence between competitors can be maintained by trace concentrations of an additional metabolite if it is taken up at sufficiently high affinity by the weaker competitor. Consistent with this prediction, targeted metabolomics and spent-medium assays identified growth-enhancing compounds at micromolar concentrations in Oa spent medium and as residues in fresh medium. Model analysis further showed that such weak positive effects can qualitatively change coexistence outcomes in chemostats while remaining undetected in standard batch interaction assays. Together, our results show that trace metabolites and subtle positive effects can reshape coexistence outcomes and should be incorporated into ecological models and interaction measurements.

## Introduction

Bacterial communities perform diverse functions, from nitrogen and carbon fixation in soil to the breakdown of complex carbohydrates in the human colon [Mylona et al., 1995, Santos Correa et al., 2023, Blaut and Clavel, 2007]. These functions depend on the interplay of multiple community members that share the same environment [Giri et al., 2019]. Predicting and controlling community function therefore requires understanding the mechanisms that allow several species to coexist. Although competition for shared resources would lead us to expect competitive exclusion to be common – especially because pairwise interaction assays often classify species pairs as competitive [MacArthur, 1970, Foster and Bell, 2012, Ono et al., 2025, Schäfer et al., 2023] – natural communities can nonetheless maintain high diversity even in nutrient-poor environments [Hardin, 1960].

Theoretical work has proposed several mechanisms that can stabilize coexistence between species that would otherwise competitively exclude one another [Chesson, 2018, Orr et al., 2025]. Recent work has shown that environmental fluctuations can stabilize communities [Letten et al., 2018, Rodríguez-Verdugo et al., 2019, Bloxham et al., 2024]: If each species is better adapted to a different environmental state, fluctuating through these states provides temporal niches, allowing them to coexist. Here, we sought to focus on a second coexistence mechanism: dependencies for essential metabolites [Shou et al., 2007, Pande et al., 2014, Kerner et al., 2012, Hammarlund et al., 2019, Hammarlund et al., 2021], whereby one species relies on another for growth. Auxotrophies – where species lack the capacity to produce all essential metabolites independently – are common in natural strains [Kehe et al., 2021, Yousif et al., 2025, Gregor et al., 2024]. If an auxotrophic species is otherwise a strong competitor, dependence on a partner can limit its growth and thereby stabilize coexistence. This logic leads to a clear prediction: if coexistence is maintained by metabolite dependency, externally supplying the missing metabolite should release the auxotroph from this constraint and allow it to exclude its competitor. This prediction was supported in an engineered *Escherichia coli*–*Salmonella enterica* mutualism, termed “feed the faster grower” [Hammarlund et al., 2019]. Because experiments in that study were performed using serial batch transfers, however, environmental fluctuations may have contributed to the outcome. Here, we tested “feed the faster grower” hypothesis using natural isolates under constant conditions.

To exclude fluctuation-based coexistence, we combined chemostat experiments and consumer– resource modeling in an auxotroph–prototroph pair originally isolated from industrial machine oils [van der Gast et al., 2004]: *Ochrobactrum anthropi*, a strong competitor on acetate that is auxotrophic for thiamine, and *Comamonas testosteroni*, a thiamine prototroph that is a weaker competitor on acetate. We first show that the two species coexist on acetate in thiamine-free chemostats, consistent with the idea that thiamine dependence stabilizes coexistence and with predictions from a consumer–resource model. However, contrary to this prediction, the species were still able to coexist when thiamine was supplied externally. We show that this persistent coexistence can be explained by either of two additional (known) mechanisms: (i) facultative cross-feeding, where metabolites released by one species boost the growth of another under strong resource limitation (e.g. [Turner et al., 1996, McCully et al., 2017]); and (ii) consumption of growth-enhancing environmental residues, where trace compounds present in the medium can benefit competitively disadvantaged species and stabilize coexistence (see discussion in [Rodríguez-Verdugo et al., 2019]). Crucially, these two mechanisms can lead to coexistence in chemostats even at very low metabolite concentrations, as long as the weaker competitor has sufficiently high affinity for them. Because such increases in growth rate have only small effects on yields in batch culture assays, interactions measured in batch cultures can miss them entirely even if they have large consequences for coexistence in continuous culture.

## Results

### A parameterized model predicts coexistence due to dependency on thiamine cross-feeding

The “feed the faster grower” hypothesis predicts that if the stronger competitor on a single limiting resource depends on a second, weaker species to survive, the two species will coexist despite competition for the limiting resource; this coexistence will be lost when the cross-fed nutrient is supplied in the environment. We tested this hypothesis (H1) in a chemostat environment with minimal environmental fluctuations using a bacterial species pair consisting of the prototroph *Comamonas testosteroni* (Ct) and the thiamine auxotroph *Ochrobactrum anthropi* (Oa), which can grow when thiamine is supplied (Fig. 1A).

**Figure 1:**
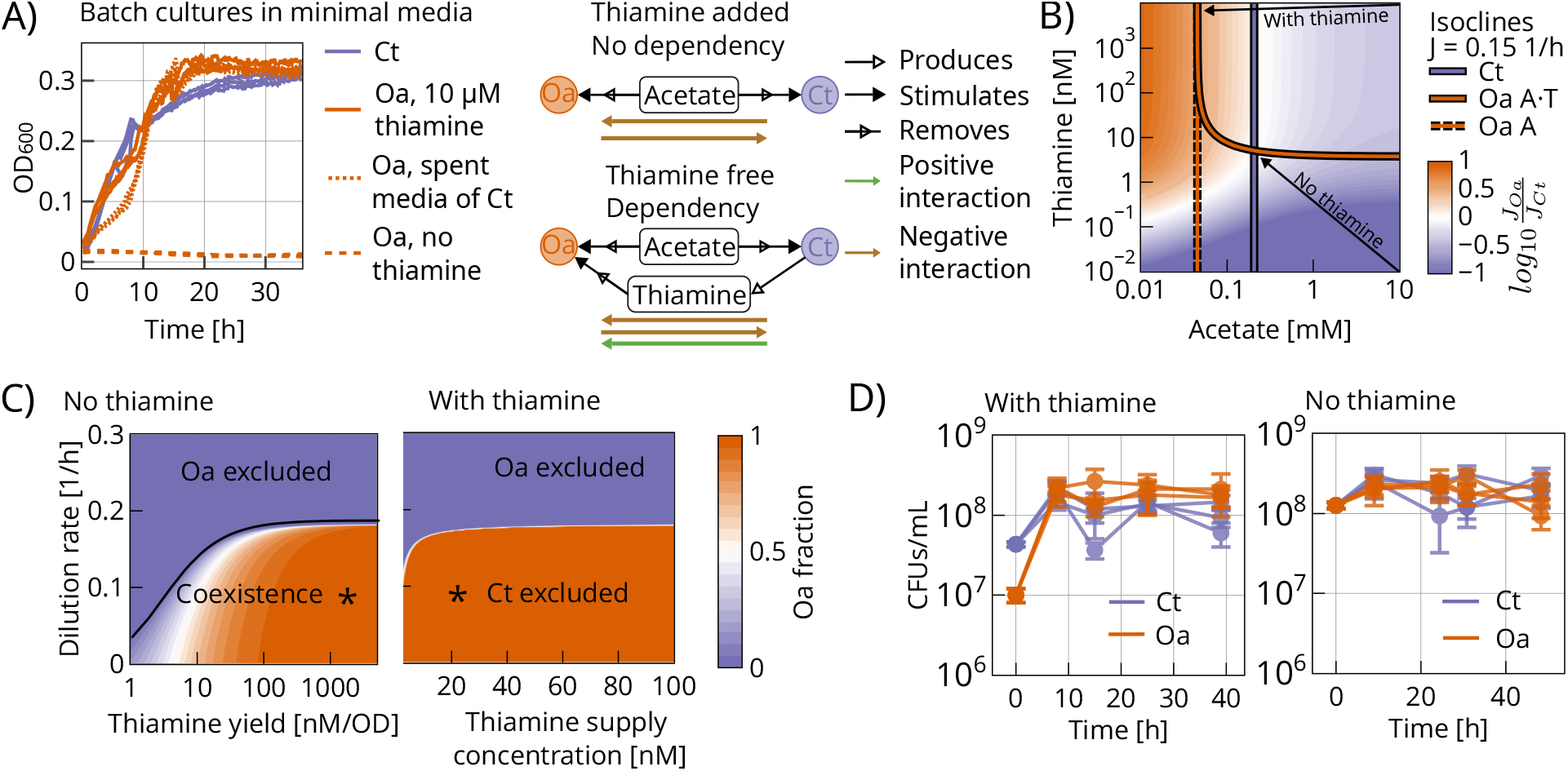
Thiamine cross-feeding explains coexistence in the model but is not required experimentally. We compare a condition where no thiamine is supplied (no thiamine), where Ct should enable Oa’s growth via thiamine cross-feeding, to a condition where thiamine is supplied in excess (with thiamine), where the two species should grow independently. Isocline analysis and chemostat simulations predict coexistence only when no thiamine is supplied, whereas chemostat experiments show coexistence in both conditions. **(A)** Left: OD_600_ of Ct (purple) and Oa (orange) in batch cultures containing minimal medium with 7.5 mM acetate (see Methods). Oa only grows if either 10 *µ*M thiamine (solid line) or the spent medium of Ct (dashed line) are added, indicating that Ct cross-feeds thiamine to Oa. Right: Inferred interaction network illustrating how thiamine modulates a metabolic dependency. In both conditions, Ct and Oa compete for acetate (bidirectional negative interaction). In the no-thiamine condition, Oa depends on Ct for thiamine, such that Ct has a positive effect on Oa’s growth. In the with-thiamine condition, this dependency is relieved, leaving acetate competition as the dominant interaction. **(B)** Isocline analysis in the acetate–thiamine space. The background shows the ratio of growth rates *J*_Oa_/*J*_Ct_ across resource concentrations. Purple and orange curves denote growth isoclines where each species reaches 0.15 h^−1^ (the dilution rate used in the experiments). For Oa, the solid orange curve shows the no-thiamine isocline, whereas the dashed orange curve shows the with-thiamine isocline. Arrows indicate simulated chemostat trajectories for inflow medium in the two conditions at *D* = 0.15 h^−1^. In the with-thiamine condition, trajectories converge to a steady state where Oa can maintain 0.15 h^−1^ at a lower acetate concentration than Ct, predicting Ct’s exclusion (R* rule). In the no-thiamine condition, Oa is co-limited by acetate and thiamine and trajectories converge to the isocline intersection where both species grow at the dilution rate and coexist. **(C)** Parameter sweeps of the chemostat model under the no-thiamine condition (left) and the with-thiamine condition (right). Left: Simulations across dilution rates and Ct thiamine production levels predict coexistence over a broad parameter range, with exclusion of Oa at high dilution rates. Thiamine yield is computed at equilibrium. * Ct reaches a low but nonzero abundance in this region (Ct = 0.00139 *OD*_600_). Right: Simulations across dilution rates and thiamine supply concentrations predict exclusion across most parameter combinations, with the excluded species depending on the dilution rate. * Ct abundance falls below the coexistence threshold (*OD*_600_ *<* 10^−6^). **(D)** Chemostat experiments with acetate as the sole limiting resource, performed in the dependency condition (no thiamine supplied; left) and the no-dependency condition (10 *µ*M thiamine supplied; right).

To test this prediction, we constructed a consumer–resource model for Ct and Oa growing in a chemostat with acetate as the sole limiting carbon source (see Methods). In the model, both species consume acetate according to Monod kinetics, while Oa’s growth additionally depends on thiamine availability. Ct releases thiamine into the environment, allowing Oa to grow in thiamine-free conditions. We parameterized the model using monoculture chemostat and batch experiments (see Methods). These measurements showed that, in the absence of thiamine limitation, Oa has a higher affinity for acetate than Ct and is therefore expected to outcompete Ct at low to intermediate dilution rates, whereas Ct should dominate at higher dilution rates because it has the higher maximum growth rate (Fig. 1C, right panel). Based on these estimates, we focused subsequent chemostat experiments on a dilution rate of *D* = 0.15 h^−1^, where the model predicts exclusion of Ct unless Oa remains limited by thiamine.

In chemostats, the limiting resource is depleted to a steady-state concentration *R*^∗^ at which per capita growth equals the dilution rate *D*; any species unable to achieve this is washed out.. We therefore analyzed the growth isoclines of Ct and Oa in acetate–thiamine space at *D* = 0.15 h^−1^ (Fig. 1B). When thiamine is not supplied, Oa is co-limited by acetate and thiamine, and its isocline intersects that of Ct, predicting coexistence. In contrast, when thiamine is supplied in excess, the Oa isocline collapses to a single acetate threshold and no longer intersects the Ct isocline. Under these conditions, Oa can maintain the dilution rate at a lower acetate concentration than Ct and is therefore predicted to exclude Ct according to the *R*^∗^ rule [Tilman, 1982].

Simulations of the full chemostat model matched the isocline analysis. In the absence of thiamine supplementation, coexistence was predicted across a broad range of Ct thiamine production levels and dilution rates. By contrast, when thiamine was supplied, exclusion was predicted across most combinations of dilution rate and thiamine concentration in the inflowing medium (Fig. 1C). Together, these results predict that coexistence should occur only when Oa remains dependent on Ct for thiamine.

### In chemostat experiments, thiamine cross-feeding is not required for coexistence

We next tested whether thiamine cross-feeding is required for coexistence in chemostat experiments. In the absence of added thiamine, Oa was unable to grow alone, both in chemostat and in batch culture (Fig. S2D, S1F), confirming that it depends on an external thiamine source under these conditions. In co-culture, Oa grew together with Ct and the two species coexisted in thiamine-free chemostats (Fig. 1D, left), consistent with the model prediction that Ct can support Oa through thiamine cross-feeding.

Unexpectedly, coexistence persisted when 10 *µ*M thiamine was added to the medium, a concentration well above that required for Oa’s growth (Fig. S1F). Under these conditions, the model predicts that the dependency should be removed and that Oa should exclude Ct. Instead, Ct remained detectable and coexisted with Oa, although at a lower abundance (Fig. 1D, right). Thus, while thiamine cross-feeding is sufficient to explain coexistence in the absence of thiamine, coexistence still occurs in conditions when thiamine cross-feeding is not required.

One possible explanation for this discrepancy is that Oa may be producing other compounds that are missing in the model. If the competitively disadvantaged species Ct is using these cross-fed compounds to grow despite acetate limitation, this could allow for coexistence (H2).

### Small concentrations of cross-fed metabolites can contribute to coexistence

To explore this possibility, we revisited the model. In thiamine-supplemented medium, the species that can maintain a growth rate equal to the dilution rate at the lowest acetate concentration should set the steady-state acetate level in the chemostat. This *R*^∗^ concentration then becomes too low for the weaker competitor to match its growth rate on acetate alone, leading to exclusion (Fig. 2A, B). Based on monoculture parameter estimates, Oa has a higher acetate affinity and is expected to dominate at the dilution rate used in our chemostat experiments (0.15h^−1^). Coexistence in the thiamine-supplemented condition at that dilution rate would then require additional compounds beyond acetate that increase Ct’s growth rate to match the dilution rate (Fig. 2C). Specifically, to account for the observed coexistence with Oa under these conditions, Ct would need an additional growth-rate contribution of 0.1 h^−1^ (Fig. 2B, C).

**Figure 2:**
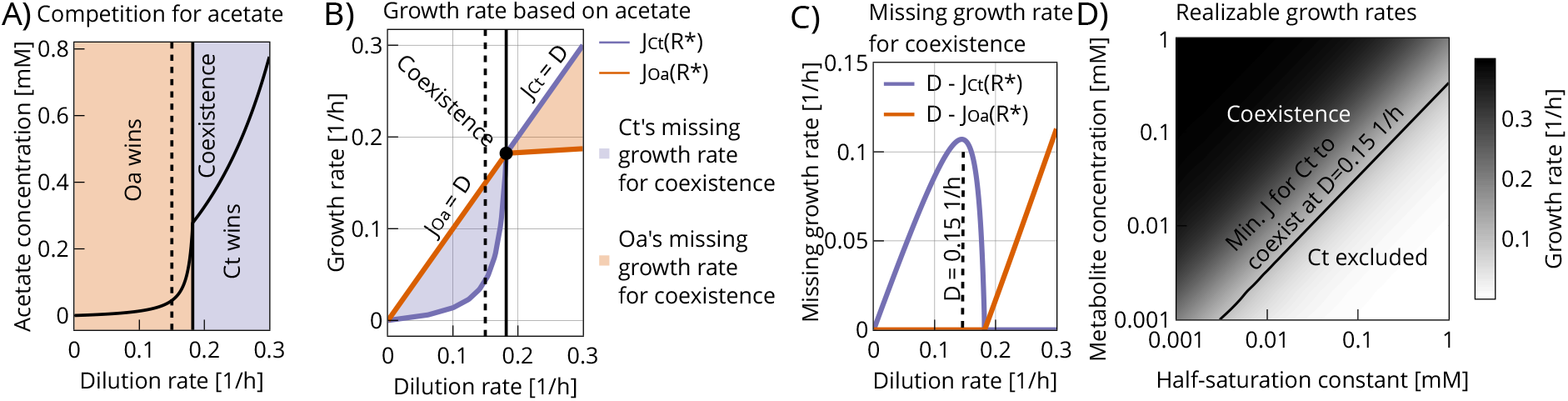
H2: In thiamine-supplemented medium, acetate competition predicts exclusion, but a small additional growth-rate contribution from a cross-fed metabolite can, in principle, stabilize coexistence. **(A)** Predicted steady-state acetate concentration (*R*^∗^) as a function of the dilution rate (*D*) under pure acetate competition. Depending on *D*, either Oa (low *D*; higher acetate affinity) or Ct (high *D*; higher maximum growth rate) sets *R*^∗^. The dashed line marks the experimental dilution rate (*D* = 0.15 h^−1^), while the solid line marks the dilution rate at which coexistence is expected. **(B)** Acetate-based growth rates of Ct and Oa evaluated at *R*^∗^(*D*) (solid curves). A species persists when its growth rate equals the dilution rate (*J* = *D*). When one species sets *R*^∗^, the partner’s acetate-based growth rate typically falls below *D* (shaded regions), implying exclusion in the absence of an additional growth contribution. **(C)** “Missing growth rate” required for coexistence, calculated by taking the difference between the two lines at each dilution rate in panel (B). This difference is the additional growth-rate contribution needed for the disadvantaged species to reach *J* = *D* at the acetate level set by the stronger competitor. At *D* = 0.15 h^−1^ (dashed line), Ct would require an additional 0.1 h^−1^ to coexist with Oa. **(D)** Growth-rate contribution from an additional metabolite to Ct’s growth as a function of metabolite concentration (*C*) and uptake affinity (*K*_*C*_) (model; Methods). The contour indicates the minimum metabolite-driven growth required for Ct to coexist with Oa at *D* = 0.15 h^−1^, illustrating that micromolar metabolite concentrations can be sufficient when uptake affinity is high.

To assess whether such a growth-rate increase could plausibly be generated by cross-feeding, we modeled growth-rate contributions from an additional metabolite as a function of its concentration and uptake affinity (half-saturation constant *K*_*m*_, see Methods). This analysis showed that even micromolar metabolite concentrations can produce substantial growth-rate contributions when uptake affinity is high (low *K*_*m*_, Fig. 2D), and that the required increase for Ct to a growth rate of 0.15 h^−1^ can, in principle, be achieved at low metabolite concentrations (Fig. 2D).

We then asked whether Oa releases growth-enhancing metabolites that are consumed by Ct. We applied targeted mass spectrometry to build profiles of metabolites in the spent chemostat medium of each species and then quantified which metabolites were depleted after growing the partner species in that spent medium in batch culture. Both species released a variety of metabolites into the chemostat medium, with many of the strongest fold changes associated with purine and pyrimidine metabolism (Fig. 3A, left column). Only a subset of released metabolites was measurably consumed by the partner (Fig. 3A, right column). The metabolite with the largest depletion by Ct was cis-aconitate, an intermediate in acetate metabolism. Cis-aconitate concentrations were highest in Oa’s spent chemostat medium (average 1.7 *µ*M) and dropped below the limit of quantification after Ct was grown in this medium (Fig. 3B), making cis-aconitate a plausible candidate cross-fed metabolite from Oa to Ct.

**Figure 3:**
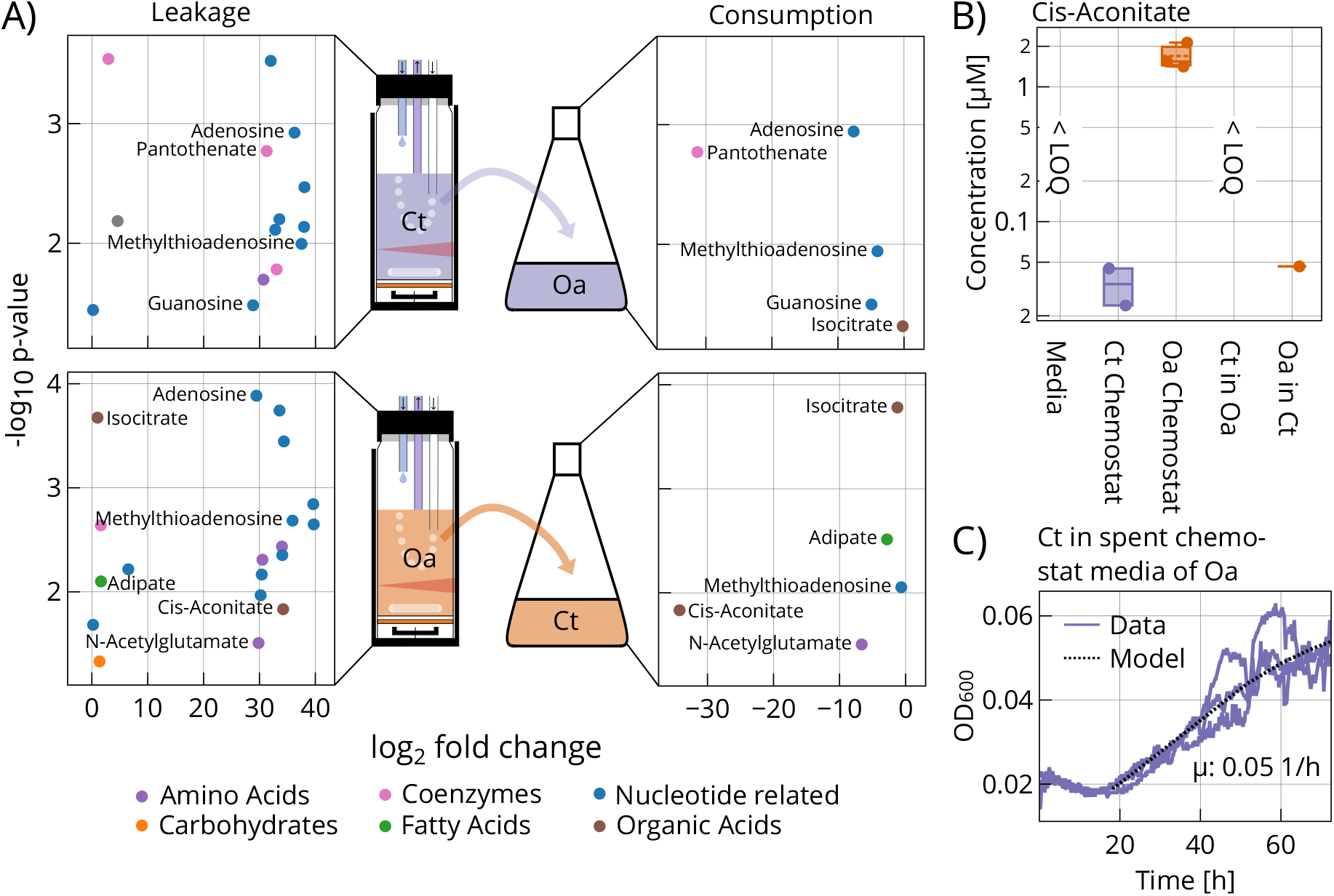
Metabolomics analyses. **(A)** Left column: Significantly increased metabolites in the spent chemostat medium of Ct (top) and Oa (bottom). Right column: Significantly depleted metabolites in the spent medium of Ct (top) and Oa (bottom) after growing them in batch in the spent chemostat medium of the other species. **(B)** Absolute quantification of cis-aconitate in all samples (LOQ: limit of quantification). Concentration of cis-aconitate was the highest in the spent chemostat medium of Oa (average 1.7 *µ*M). Concentration below LOQ when Ct grown in the spent chemostat medium of Oa. **(C)** Growth curve of Ct grown in batch in the spent chemostat media of Oa.

However, when we grew Ct in Oa’s spent chemostat medium in batch, we observed only slight growth, with a maximum growth rate (0.05 h^−1^), below the required increase of 0.1 h^−1^ estimated from the chemostat analysis (Fig. 3C). Interestingly, Ct also showed growth in its own spent chemostat medium, whereas Oa showed no growth in either its own or Ct’s spent medium (Fig. S2A-C). Together, these results indicate metabolite exchange occurs from Oa to Ct, but the growth benefit detectable in batch culture is small relative to the increase required to explain coexistence at 0.15 h^−1^.

### Residues in growth medium can contribute to coexistence independently of species–species interactions

We next asked what else could explain the growth increase needed for coexistence. Our next hypothesis (H3) is that the environment itself might contain low concentrations of metabolites contaminating the growth medium. To test this hypothesis, we first quantified trace metabolites in fresh medium (7.5 mM acetate, 0.01 *µ*M thiamine) using mass spectrometry. Six metabolites were above the detection level (Fig. S4A). While peak areas are reported in arbitrary units, they indicate that these compounds are present in fresh medium and reproducibly detectable. Absolute quantification of selected metabolites revealed citrate at an average concentration of 15.87 *µ*M and lactate at 7.6 *µ*M, along with lower concentrations of isoleucine and glutamine (Fig. S4B).

Having found that growth media can contain unwanted carbon sources, we tested whether even the minimal medium lacking any carbon source can support growth in continuous culture. We grew Ct and Oa in co-culture at a dilution rate of 0.15 h^−1^ in this carbon-free medium. After an initial decline of both species, their populations stabilized after 40 h and coexisted in the chemostat for the rest of the experiment (Fig. 4A). This supports H3 and shows that chemical impurities in growth medium ingredients can contribute to growth independently of species–species interactions.

**Figure 4:**
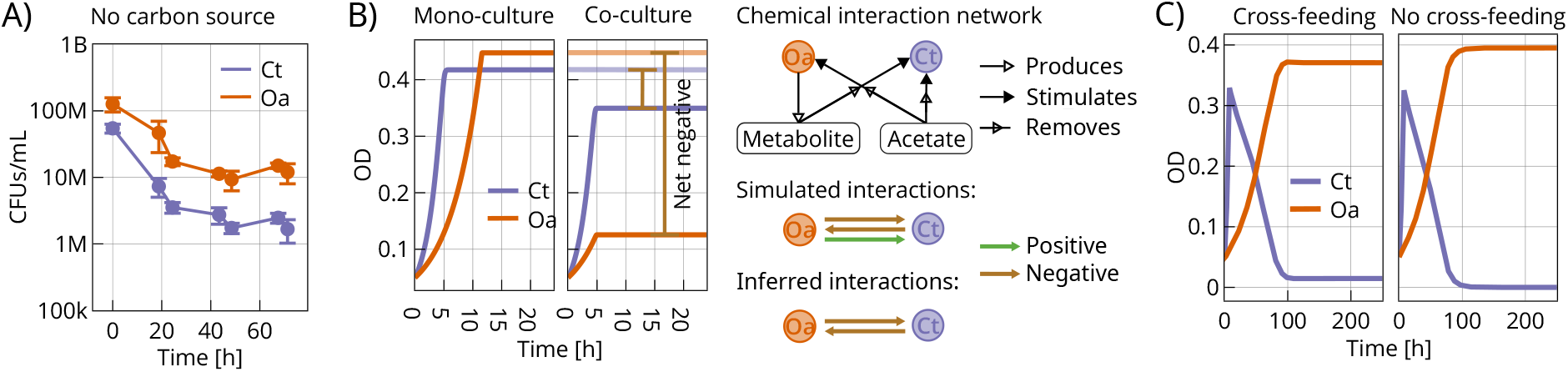
No-carbon chemostat experiment and batch/chemostat simulations illustrating that acetate competition can mask positive interactions in batch cultures. **(A)** Chemostat co-culture of Ct and Oa in minimal medium without added carbon source at *D* = 0.15 h^−1^. After an initial decline, both populations stabilized by ∼ 40 h and persisted together until the end of the experiment. **(B)** Batch-culture simulations parameterized with Ct and Oa monoculture estimates and including an additional positive interaction from Oa to Ct. Left: Simulated growth curves for Ct and Oa in mono- and co-culture (thiamine supplied). Right: Schematic of the modeled interaction network. Both species consume acetate (competition), and Oa additionally releases a low-concentration metabolite (total production 200 *µ*M) that Ct can consume. Despite this facilitation, co-culture yields are reduced relative to monoculture due to acetate competition, so the interaction would be classified as net negative based on yield. **(C)** The same positive interaction can qualitatively change outcomes in chemostat simulations at *D* = 0.15 h^−1^. Right: Without cross-feeding, Oa excludes Ct. Left: With the additional Oa→Ct cross-feeding interaction, Ct and Oa coexist.

### Competition masks positive interactions when quantified in batch cultures

Our results indicate that very small concentrations of growth-enhancing compounds, whether they are produced by a partner species (H2) or present in the abiotic environment (H3) can increase growth rates in chemostats sufficiently to enable coexistence. In the case of cross-feeding, we wondered why we were not able to detect this positive interaction in standard batch assays, which are the gold standard for measuring interactions [Mitri and Foster, 2013]. In these assays, interactions are typically classified by comparing yields between mono- and co-cultures. We reasoned that this approach will surely reveal competitive interactions that can strongly reduce yields in co-culture, but may mask facilitation through the production of metabolites at low concentrations that only have small positive effects on yields.

To illustrate this point, we simulated mono- and co-culture batch cultures using the Ct and Oa parameter estimates (Fig. 4B). In the co-culture simulations, we included a weak positive interaction in which Oa releases a metabolite at low concentrations that Ct can consume in addition to acetate (see Supporting Information for the model description; Fig. 4B; Fig. S5). Despite this facilitation, competition for acetate reduced the yield of both species relative to the monoculture, a pattern that would typically be interpreted as a net negative interaction. In these batch simulations, the positive effect on Ct was therefore masked by the large impact of resource competition on yield. In contrast, simulating the same interaction in a chemostat at a dilution rate of 0.15 h^−1^ showed that despite being mediated by small concentrations, the resulting increase in growth rate can be significant, qualitatively changing community outcomes: with the positive interaction Ct coexists with Oa, whereas without it Oa excludes Ct (Fig. 4C). Together, these simulations highlight that interaction screens based on batch yield can miss facilitation even when it is sufficient to stabilize coexistence in continuous culture.

## Discussion

A central challenge in microbial ecology is reconciling the expectation of competitive exclusion under shared resource limitation with the high diversity observed in nature. Using a tractable auxotroph–prototroph pair (Ct and Oa) in acetate-limited chemostats, we evaluated three candidate mechanisms that can counteract exclusion: (H1) an obligate metabolic dependency, (H2) facultative cross-feeding that provides an increase in growth rate to the disadvantaged species, and (H3) an increase in growth rate arising from trace, growth-enhancing residues present in the environment. Together, our results show that coexistence under strong resource limitation can be stabilized by multiple, potentially overlapping effects, including effects that have little impact on batch yield.

The analysis of H1, the “feed the faster grower” hypothesis, illustrates how an obligate dependency can stabilize coexistence in a chemostat: when Oa depends on Ct for thiamine, it cannot exclude Ct even when it is the stronger competitor for acetate. This feedback is readily captured by consumer–resource theory and is reflected in our isocline analysis and model simulations. Such dependency-based feedbacks are likely relevant beyond this system because vitamin and amino-acid auxotrophies are common in bacteria [Kaeberlein et al., 2002, Embree et al., 2015], and comparative analyses suggest that metabolic dependencies are widespread across environments [Kehe et al., 2021, Zelezniak et al., 2015]. However, a key experimental result was that coexistence persisted when we relieved the dependency on the prototroph by supplying thiamine. This discrepancy suggests a broader point: while dependency-based feedback is sufficient for coexistence in theory, it may be difficult to isolate experimentally because additional interactions can act in parallel. This contrasts with systems in which interactions can be tuned more directly by supplementing exchanged metabolites, for example in engineered amino-acid auxotroph pairs [Hoek et al., 2016]. This highlights the importance of building on the work done with genetically engineered model organisms and studying interactions between natural isolates, where the entire metabolic architecture of interacting partners can differ.. Our results therefore emphasize that interpreting coexistence outcomes solely through the lens of a focal dependency can be misleading if other growth-enhancing compounds are present.

When we investigated facultative cross-feeding (H2), we found that low concentrations of growth-enhancing metabolites can be sufficient to promote coexistence under acetate competition. The magnitude of the growth-rate increase that a disadvantaged species can obtain from an additional metabolite depends strongly on its uptake affinity: higher affinities (lower half-saturation constants) allow substantial growth-rate gains even at micromolar concentrations. However, half-saturation constants are poorly constrained for many metabolites and species. A literature compilation [Fink et al., 2023] reports values spanning orders of magnitude (Fig. S3), but many fall in the micromolar range, consistent with appreciable growth-rate contributions at low concentrations. We also found that the additional growth-rate contribution required for coexistence depends on the dilution rate and on which species is competitively disadvantaged. In particular, the required increase is smallest near batch-like conditions and near the dilution rate at which the two species have similar acetate-based growth rates, implying that weak positive effects can be sufficient over parts of the dilution-rate range. Mechanistically, our metabolomics analysis identified cis-aconitate as a candidate metabolite cross-fed from Oa to Ct at *D* = 0.15 h^−1^. Cis-aconitate is a TCA-cycle intermediate linked to acetate metabolism [Cozzone, 1998, Petushkova et al., 2021], and uptake of such intermediates could provide a small but consequential growth-rate contribution under acetate limitation. At the same time, the modest growth observed in spent-medium assays suggests that survival of Ct in the chemostat may depend on multiple low-concentration compounds, on continuous production and consumption in chemostats, or on effects that influence growth rate more strongly than endpoint yield.

Our H3 results further suggest that growth-enhancing compounds need not originate from an interaction partner. Freshly prepared medium can contain trace organic compounds introduced with chemical reagents, and we detected micromolar concentrations of citrate and lactate in our medium (Fig. S4B). Trace residues at the tens-of-micromolar level are also plausible from reagent purity alone; for example, using 7.5 mM acetate of ≥ 99% purity implies potentially up to 1% of the carbon source as uncharacterised impurities of unknown bioavailability. Even if individual compounds are not the dominant carbon source, such residues can contribute growth-rate increments that matter under strong acetate limitation, particularly when uptake affinities are high. Consistent with this possibility, in chemostats without added carbon source we observed an initial decline followed by stabilization and persistence of both Ct and Oa. While this does not uniquely identify the underlying carbon source(s), it supports the broader conclusion that trace residues can influence dynamics in continuous culture and should be considered when interpreting coexistence outcomes in defined media.

A practical implication of these findings is methodological. Pairwise interactions are often inferred from batch co-culture yields, an approach that emphasizes the strong yield-reducing effects of resource competition. Our simulations highlight that weak facilitation mediated by low-concentration metabolites can be masked in batch yields, even when the same facilitation is sufficient to stabilize coexistence in chemostats. This suggests that interaction summaries derived from batch assays may underestimate weak positive effects [Foster and Bell, 2012, Palmer and Foster, 2022, Meacock and Mitri, 2025] and motivates complementary measurements that are sensitive to small growth-rate contributions under nutrient limitation.

Our study has several limitations. First, both the model and its parameterization simplify metabolism into a small number of resources and growth terms; real metabolite exchange is likely multi-dimensional, with mixtures of compounds contributing to growth. Second, experimental chemostats are not perfectly mixed and biofilm formation can provide refuges or inocula that complicate interpretations of exclusion dynamics. Third, our ability to link specific metabolites to quantitative growth-rate contributions is constrained by detection limits, incomplete coverage of the metabolite space, and uncertainty in uptake affinities. Finally, practical constraints limit chemostat run times; although steady states were reached in our experiments, longer runs would further strengthen inferences about stability. Future work that directly measures uptake kinetics for candidate metabolites, quantifies fluxes, and manipulates residue levels in defined media would strengthen causal inference.

Overall, our results show that positive effects that are weak by standard batch-yield criteria can nonetheless be decisive for coexistence in continuous culture. This supports the need to incorporate weak facilitation and environmental residues into ecological models and to develop experimental approaches that quantify interactions in ways that reflect the dynamical conditions under which coexistence is maintained. In nutrient-limited environments, where growth can be sensitive to trace compounds, such weak effects may represent an underappreciated but pervasive contributor to microbial diversity.

## Materials and methods

### Bacterial strains and growth medium

We used Comamonas testosteroni (Ct) and Ochrobactrum anthropi (Oa). These isolates were kindly provided by Christopher van der Gast and Ian Thompson and were originally isolated from industrial wastewater, as previously described [Piccardi et al., 2019]. Assembled genomes are available under BioProject accession PRJNA991498.

Minimal medium was prepared from 10× M9, 50× HMB, 10× potassium acetate, and, when indicated, 10× thiamine. Media were brought to 1 L with water, adjusted to pH 7.4, and filter-sterilized through 0.22 *µ*m filters. Recipes for 10× M9 and HMB were as described previously [Dos Santos et al., 2022]. The 10× potassium acetate stock (75 mM) was prepared by dissolving 7.36 g potassium acetate in 1 L water and filter-sterilizing. The 10× thiamine stock (0.1 mM thiamine hydrochloride) was prepared by dissolving 33.7 mg thiamine hydrochloride in 1 L water and filter-sterilizing. Final media contained 7.5 mM acetate and, when supplemented, 10 *µ*M thiamine.

### Culture conditions

Single colonies of Ct and Oa were inoculated into 50 mL Erlenmeyer flasks containing acetate minimal medium supplemented with thiamine and incubated at 28 °C with shaking at 200 rpm. Ct precultures were grown for 24 h and Oa precultures for 48 h. Overnight cultures were then started at OD_600_ = 0.05 on the day before each experiment and used as inocula unless stated otherwise.

### Chemostat setup

Chemostat experiments were performed using a modified version of the Chi.Bio platform[Steel et al., 2020]. Cultures were grown in 30 mL glass vials sealed with 20 mm septa and fitted with PTFE tubing for medium inflow, outflow, sampling, and aeration or pressure release. Cultures were mixed with a magnetic stir bar and maintained at 28 °C.

Fresh medium was supplied from a reservoir using an Ismatec IPC multichannel peristaltic pump. Effluent was removed through a separate outflow line, the height of which was adjusted to maintain a working volume of 20 mL. Cultures were aerated using an aquarium air pump connected through an air filter, with one port left open for pressure release. Assembled reactors were autoclaved before use. Experiments were initiated by filling the reservoir with sterile medium and inoculating the reactor with 20 mL of medium containing the appropriate inoculum. Samples of 700 *µ*L were withdrawn through the septum using a syringe. Optical density was monitored using the built-in Chi.Bio laser sensor and converted to OD_600_ using calibration curves (Fig. S6). Raw Chi.Bio laser sensor values were converted to OD_600_ using a two-point calibration curve (Fig. S6). Two samples were measured both with the Chi.Bio sensor and independently by spectrophotometry, and the resulting paired values were used to determine the slope *k* and intercept *b* of the calibration equation *OD*_600_ = *k* log_10_(*R*) + *b*.

### Monoculture and community chemostat experiments

For monoculture chemostat experiments, Ct and Oa were each grown in triplicate in thiamine-supplemented minimal medium. Chemostats were inoculated at OD_600_ = 0.05. At the end of each experiment, 20 mL culture was harvested by centrifugation at 4000 rpm for 10 min, and the supernatant was filter-sterilized twice. Filtrates were stored at −80 °C for targeted mass spectrometry. In addition, 10 mL of reservoir medium was collected and stored at −80 °C.

For community chemostat experiments, Ct and Oa were grown together in triplicate in minimal medium with or without thiamine supplementation. Communities were inoculated at OD_600_ 0.1. Species abundances were quantified by CFU counting. Chemostat samples were serially diluted to 10^−7^ in 96-well plates, and 5 *µ*L of each dilution was spotted onto tryptic soy agar (TSA) plates and TSA supplemented with colistin (10 *µ*g/mL). Ct colonies were counted on TSA after 24 h incubation at 28 °C. Oa, which is naturally resistant to colistin, was counted on colistin plates after 48 h.

### Spent-medium rescue and thiamine gradient assays

For the spent-medium rescue assay (Fig. 1A) and the thiamine gradient experiment (Fig. S1F), thiamine-starved Oa inoculum was prepared by growing Oa for 48 h in a chemostat supplied with thiamine-free minimal medium. Cells were harvested from 8 mL culture by centrifugation at 4000 rpm for 10 min, resuspended in thiamine-free minimal medium, and adjusted to OD_600_ = 0.05.

To generate Ct spent medium for the rescue assay, Ct was grown to stationary phase in thiamine-free minimal medium. Cultures were centrifuged at 4000 rpm for 10 min, and the supernatant was filter-sterilized twice through 0.22 *µ*m filters. Filtered Ct spent medium (5 mL) was mixed 1:1 with minimal medium containing 15 mM acetate, resulting in a final acetate concentration of 7.5 mM. For plate-reader assays, 180 *µ*L of this medium was inoculated with 20 *µ*L of OD-adjusted, thiamine-starved Oa culture.

For the thiamine gradient experiment, a 1:10 dilution series of thiamine was prepared in thiamine-free minimal medium in a 96-well plate, spanning final concentrations from 10 *µ*M to 0.01 nM, together with a no-thiamine control. Wells were inoculated with 20 *µ*L of thiamine-starved Oa inoculum to a final volume of 200 *µ*L. Growth curves were measured in triplicate in a BioTek Synergy H1 plate reader with orbital shaking at 28 °C, recording OD_600_.

### Growth in spent chemostat medium

Ct and Oa were grown in their own or the partner species’ spent chemostat medium collected from monoculture chemostat experiments. Before use, spent medium was filtered twice through 0.22 *µ*m filters. Overnight cultures were harvested by centrifugation (8 mL, 4000 rpm, 10 min), washed three times with 1 mL PBS, and adjusted to OD_600_ = 0.05. Spent medium (180 *µ*L) was inoculated with 20 *µ*L washed culture, and growth curves were measured in triplicate in a BioTek Synergy H1 plate reader with orbital shaking at 28 °C, recording OD_600_. In parallel, 5 mL cultures were grown in the corresponding spent media for 72 h, centrifuged at 4000 rpm for 10 min, and filter-sterilized twice. Filtrates were stored at −80 °C for targeted mass spectrometry.

### Targeted mass spectrometry

For targeted LC–MS/MS analysis, 500 *µ*L of each sample was submitted to the UNIL Metabolomics Facility using the high-coverage targeted analysis of polar metabolites workflow. Briefly, 20 *µ*L of sample was extracted with 80 *µ*L ice-cold methanol, vortexed for 30 s, and centrifuged for 15 min at 15000 rpm and 4 °C. Supernatants were analyzed by hydrophilic interaction chromatography coupled to tandem mass spectrometry on an Agilent 6496 iFunnel instrument using multiple reaction monitoring in positive and negative ionization modes. Analytical conditions were as described previously [van der Velpen et al., 2019, Gallart-Ayala et al., 2018]. Raw data were processed using MassHunter Quantitative Analysis (Agilent Technologies). Peak areas were used for relative comparisons, with drift correction based on pooled quality-control samples. Absolute quantification of selected metabolites was performed using stable isotope-labeled internal standards and calibration curves. Peak integration was manually curated, and concentrations were reported within the linear range of the corresponding calibration curves.

### Mathematical model

We developed a Monod-type chemostat model describing Ct and Oa growth on acetate. Under thiamine-free conditions, Oa growth additionally depended on thiamine produced by Ct, whereas under thiamine supplementation Oa growth was assumed to be effectively independent of thiamine. To test whether weak positive interactions could stabilize coexistence despite acetate competition, we extended the model by including an additional metabolite produced by Oa and consumed by Ct. Model parameters were constrained using monoculture chemostat OD_600_ trajectories and batch growth assays. Maximum growth rates were estimated from sliding-window regressions of log-transformed OD_600_ trajectories, acetate Monod constants were inferred by least-squares fitting of monoculture chemostat dynamics, and the thiamine Monod constant of Oa was fitted using batch growth data from the 1 nM thiamine condition. Parameters not directly constrained by the data, including Ct growth on the additional metabolite and the corresponding production term, were set manually for illustrative simulations. Full model equations, parameter values, and fitting details are provided in Supplementary Note S1 and Table T1.

## Data and code availability

All the data is exported in a single excel file [here](https://github.com/nahanoo/ChiBioFlow/blob/main/data/data.xlsx). Code is available [here](https://github.com/nahanoo/ChiBioFlow/tree/main/scripts).

## Acknowledgments

We thank the Metabolomics and Lipidomics team at the Faculty of Biology and Medicine, University of Lausanne, Switzerland, for metabolomic/lipidomic/metabolite analysis. We thank all members of the Mitri lab at the University of Lausanne for discussions and feedback.

