## Supporting Figures for "Micromolar concentrations of metabolites enable coexistence of bacterial species in chemostats"

397 # Supporting figures

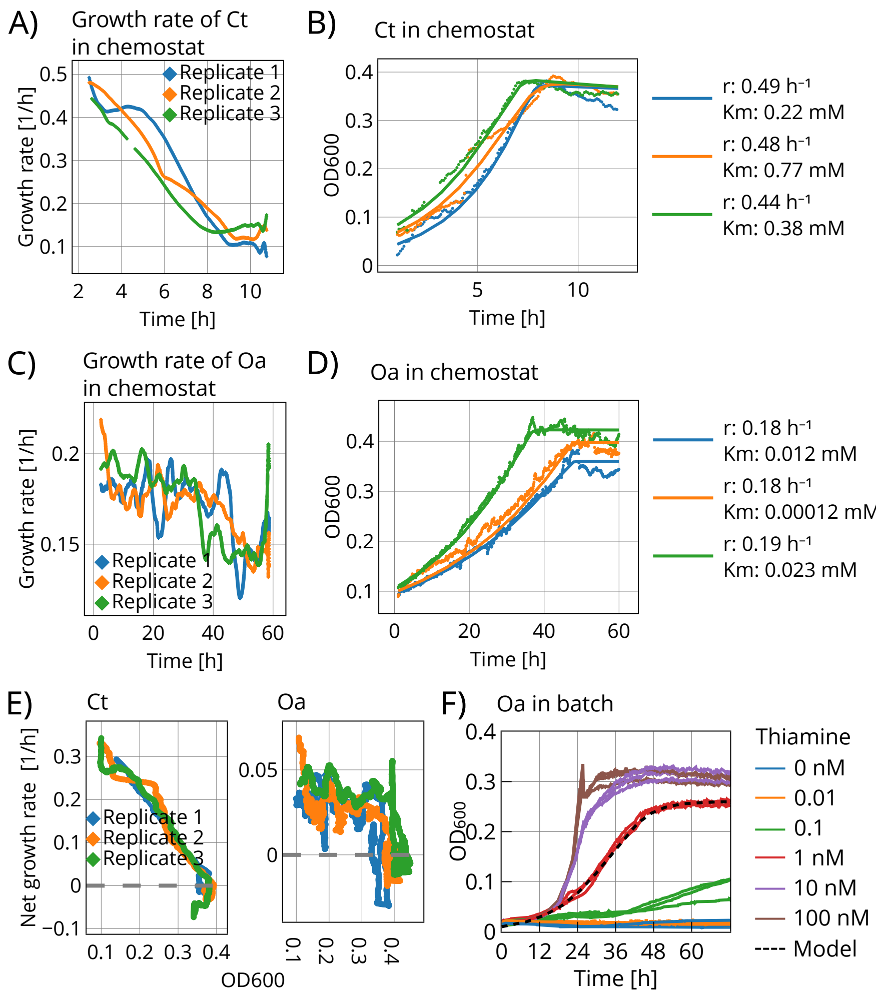


Figure S1: Ct and Oa chemostat and batch monoculture experiments used to infer growth parameters. **(A–C)** Instantaneous growth rates of Ct and Oa in chemostat monocultures, estimated as the slopes of linear fits to log-transformed OD within sliding windows (window size: 5 h). **(B–D)** OD of Ct and Oa in chemostat monoculture experiments (dots) and corresponding simulations (solid lines) using the maximum instantaneous growth rates and $K_A$, obtained by least-squares fitting of a consumer–resource model. **(E)** Instantaneous net growth rates of Ct and Oa in chemostats plotted as a function of $OD_{600}$. **(F)** Thiamine-starved Oa from the chemostat experiment with no thiamine (S2 panel D) was grown in batch culture with different thiamine concentrations. Concentrations $\geq 100$ nM did not result in higher growth rates or yields and are therefore not shown. Affinity of Oa to thiamine was inferred by fitting $K_{\text{Oa},T}$ to the growth curve at 1 nM thiamine (dashed line) obtained by least-squares fitting of a consumer–resource model.

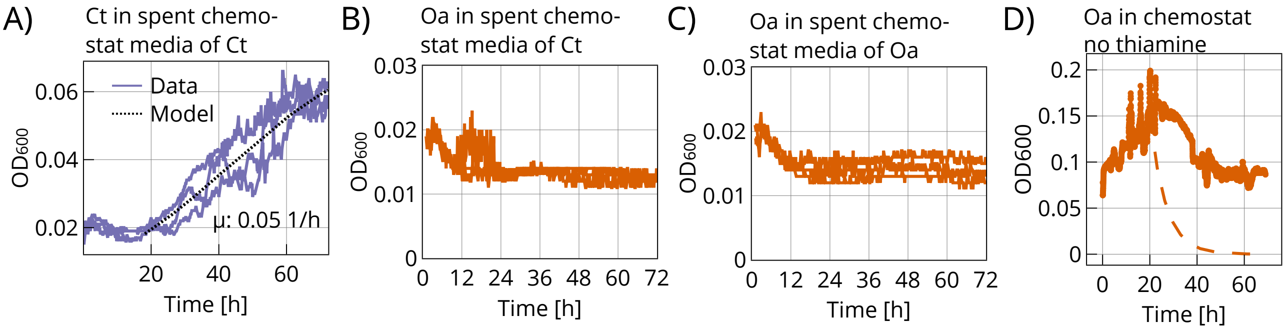


Figure S2: **(A)** Batch growth of Ct in spent medium collected from a Ct chemostat. The dashed line indicates the logistic growth model fitted by least squares, from which the maximum growth rate $\mu$ was inferred. **(B–C)** Oa grown in batch in the spent chemostat medium of Ct (B); Oa grown in bath in spent chemostat medium of Oa (C). **(D)** Chemostat mono-culture experiment with thiamine-starved Oa in minimal medium containing 7.5 mM acetate (no thiamine). Oa showed a small initial increase in OD in the chemostat, likely due to thiamine carryover, but did not sustain growth because of its thiamine auxotrophy, as confirmed by the absence of growth in subsequent batch assays. A low residual OD signal persisted in the reactor over time. The dashed line shows the simulated no-growth washout trajectory starting at 20 h.

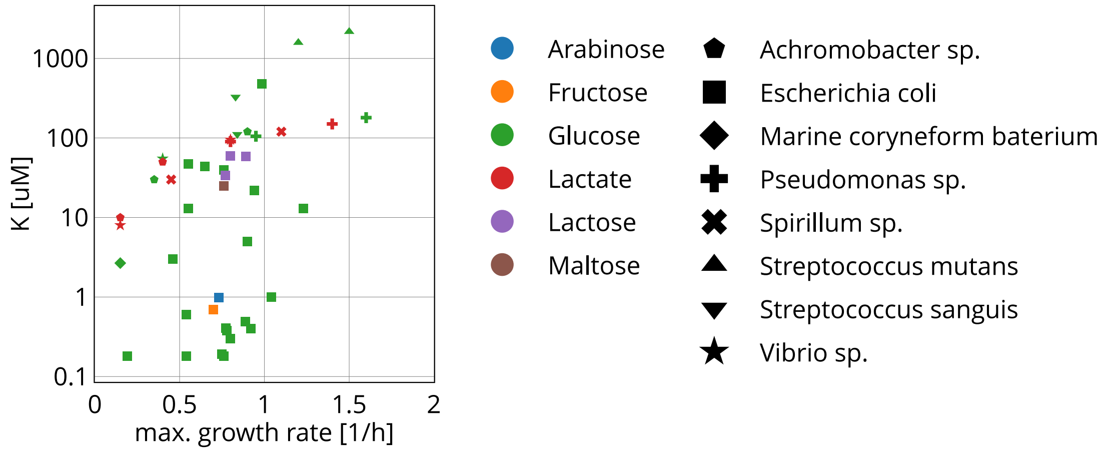


Figure S3: Half-saturation constant $K$ versus maximum growth rates of different bacteria for different carbon sources (adapted from Fink et al. [Fink et al., 2023]).

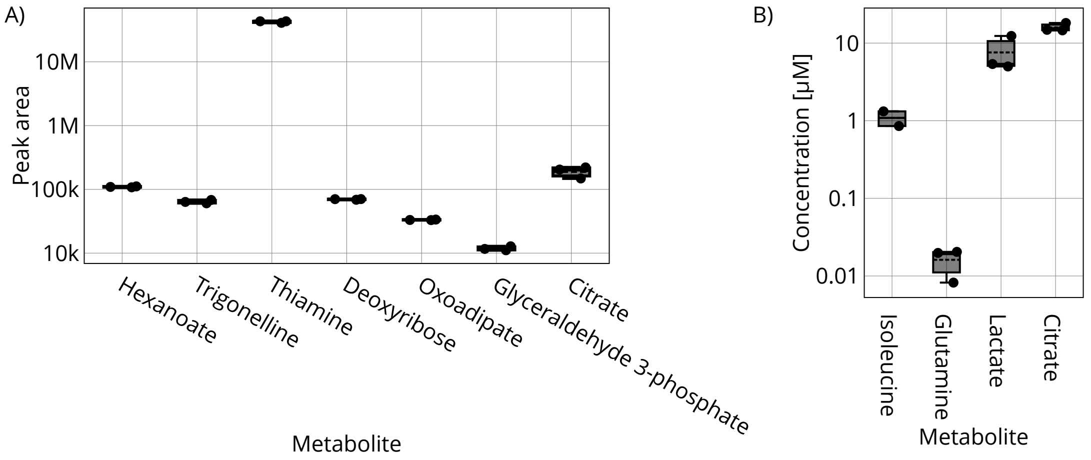


Figure S4: **(A)** Peak areas of different metabolites present in fresh minimal medium containing 7.5 mM acetate and 10 $\mu$M thiamine. Identified metabolites are likely due to chemical contamination. Note that peak values are not comparable between different metabolites. **(B)** Absolute quantification of targeted metabolites (against standard curves) in fresh minimal media containing 7.5 mM acetate and 10 $\mu$M thiamine.

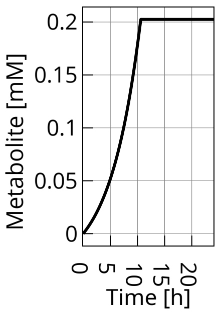


Figure S5: Concentration of produced metabolite by Oa in simulated batch mono-culture. The same parameters used for this simulation were used to simulate the co-culture batch experiments and the cross-feeding chemostat experiment.

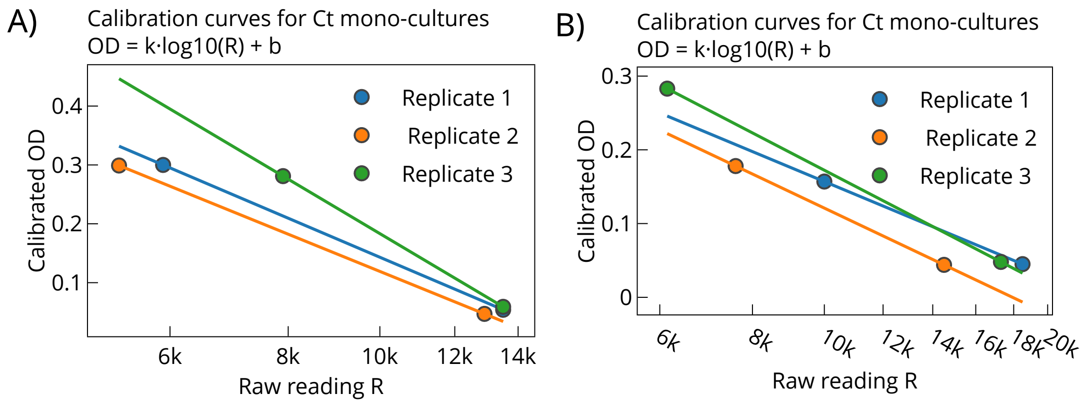


Figure S6: Calibration curves used to convert the raw Chi.Bio signal to $OD_{600}$. Two reference samples were measured with a spectrophotometer and used to determine the calibration slope ($k$) and intercept ($b$) for OD calculation. **(A)** Calibration curves measured for Ct mono-culture experiments. **(B)** Calibration curves measured for Oa mono-culture experiments.
