## Supplementary Note for "Micromolar concentrations of metabolites enable coexistence of bacterial species in chemostats"

### Supplementary Note S1: Mathematical model

#### Growth rates

We modeled the growth rates $J$ of Ct and Oa in the acetate–thiamine resource space using Monod kinetics [Monod, 1978]. For Oa, we assume acetate and thiamine are essential (interactive) resources by multiplying Monod terms for acetate and thiamine [Egli et al., 1993, Tilman, 1980]. Let $A$ denote acetate concentration, $T$ thiamine concentration, $r$ the maximum growth rate, and $K$ the Monod constant. The growth functions are:

$$J_{Ct}(A) = r_{Ct,A} \frac{A}{A + K_{Ct,A}} \tag{E1}$$

$$J_{Oa}(A, T) = r_{Oa,A,T} \frac{A}{A + K_{Oa,A}} \frac{T}{T + K_{Oa,T}} \tag{E2}$$

#### Thiamine cross-feeding

We next describe the chemostat model used to simulate a two-species community with thiamine cross-feeding. Both species consume acetate and convert acetate into biomass with yields $Y_{Ct,A}$ and $Y_{Oa,A}$, respectively. Acetate is supplied from a reservoir at concentration $M_A$ and removed from the vessel (together with cells) at dilution rate $D$. Acetate dynamics are:

$$\frac{dA}{dt} = D(M_A - A) - J_{Ct} \frac{Ct}{Y_{Ct,A}} - J_{Oa} \frac{Oa}{Y_{Oa,A}} \tag{E3}$$

Population dynamics follow:

$$\frac{dCt}{dt} = Ct \left( J_{Ct} - D \right) \tag{E4}$$

$$\frac{dOa}{dt} = Oa \left( J_{Oa} - D \right) \tag{E5}$$

We assume that thiamine is produced by Ct during growth. With biomass measured in OD, growth rates in h$^{-1}$, and thiamine concentration $T$ in nM, a production coefficient $q_{Ct,T}$ in nM/OD converts Ct biomass production $J_{Ct}Ct$ from OD/h to a thiamine production flux in nM/h. Thiamine consumption by Oa is described by a yield $Y_{Oa,T}$ in OD/nM, such that $J_{Oa}Oa/Y_{Oa,T}$ also has units nM/h. Thiamine is additionally washed out at dilution rate $D$ in h$^{-1}$.

$$\frac{dT}{dt} = q_{Ct,T} J_{Ct} Ct - J_{Oa} \frac{Oa}{Y_{Oa,T}} - DT \tag{E6}$$

#### Thiamine supplementation

When simulating the community with thiamine in the inflowing medium, we assume thiamine is supplied in large excess relative to $K_{Oa,T}$, so Oa growth is effectively independent of $T$. We therefore simplify:

$$J_{Oa}(A) = r_{Oa,A} \frac{A}{A + K_{Oa,A}} \tag{E7}$$

Acetate dynamics remain as in Eq. E3 with the updated $J_{Oa}$. Thiamine production by Ct is neglected under supplementation, and thiamine dynamics reduce to:

$$\frac{dT}{dt} = D(M_T - T) - J_{Oa} \frac{Oa}{Y_{Oa,T}} \tag{E8}$$

Growth of Ct and Oa is modeled as in Eqs. E4 and E5 with $J_{Oa}$ given by Eq. E7. For Fig. 1C (dependency condition), we varied $q_{Ct,T}$ and reported steady-state outcomes.

#### Realizable growth rates on growth-enhancing metabolites

To generate Fig. 2D, we computed growth-rate contributions from an additional metabolite with Monod kinetics, $J(C) = r\,\frac{C}{C+K}$, over metabolite concentrations $C \in [10^{-3}, 1]$ mM and Monod constants $K \in [10^{-3}, 1]$ mM, using $r = 0.4$ h$^{-1}$.

#### Oa cross-feeds a metabolite to Ct

To illustrate how weak cross-feeding can enable coexistence while remaining masked in batch yield assays, we extended the chemostat model by adding a metabolite $C$ produced by Oa and consumed by Ct. Biomass densities are measured in OD, growth rates in h$^{-1}$, and metabolite concentration $C$ in mM. We assume that metabolite production is proportional to Oa growth, with production coefficient $q_{Oa,C}$ in mM/OD, such that the production term $q_{Oa,C} J_{Oa} Oa$ has units mM/h. Ct consumes $C$ with Monod kinetics.

$$\frac{dC}{dt} = q_{Oa,C} J_{Oa} Oa - r_{Ct,C} \frac{C}{C + K_{Ct,C}} Ct - DC \tag{E9}$$

Acetate dynamics remain as in Eq. E3. Ct can additionally grow on $C$, so its dynamics become:

$$\frac{dCt}{dt} = Ct\left(J_{Ct} + r_{Ct,C}\frac{C}{C + K_{Ct,C}} - D\right) \quad \text{(E10)}$$

### Parameterization of the model

Table T1 lists parameter values used in the models above.

| Parameter | Description | Value | Unit |
|---|---|---|---|
| $r_{Ct,A}$ | Max.\growth rate of \textit{Ct} on acetate | 0.47 | 1/h |
| $r_{Oa,A,T}$ | Max.\growth rate of \textit{Oa} on acetate and thiamine | 0.19 | 1/h |
| $r_{Ct,C}$ | Max.\growth rate of \textit{Ct} on metabolite $C$ | 0.47 | 1/h |
| $K_{Ct,A}$ | Monod constant of \textit{Ct} for acetate | 0.44 | mM |
| $K_{Ct,C}$ | Monod constant of \textit{Ct} for metabolite $C$ | 0.44 | mM |
| $K_{Oa,A}$ | Monod constant of \textit{Oa} for acetate | 0.012 | mM |
| $K_{Oa,T}$ | Monod constant of \textit{Oa} for thiamine | 1 | nM |
| $Y_{Ct,A}$ | Yield of \textit{Ct} on acetate | 0.049 | OD/mM |
| $Y_{Oa,A}$ | Yield of \textit{Oa} on acetate | 0.053 | OD/mM |
| $Y_{Oa,T}$ | Yield of \textit{Oa} on thiamine | 0.25 | OD/nM |
| $q_{Ct,T}$ | Thiamine production coefficient of \textit{Ct} | 448.4 | nM/OD |
| $q_{Oa,C}$ | Production coefficient of \textit{Oa} for metabolite $C$ | 18.87 | mM/OD |
| $D$ | Dilution rate | 0.15 | 1/h |

Table T1: Parameters and values used in the model. Parameters highlighted in yellow were obtained by fitting the model to experimental data. All other parameters were set manually, except for the dilution rate $D$, which was controlled experimentally.

#### Maximum growth rates

The maximum growth-rate parameters $r_{Ct,A}$ and $r_{Oa,A,T}$ were estimated from monoculture chemostat $OD_{600}$ trajectories (SFig. S1B-D). For each replicate, instantaneous growth rates were calculated by linear regression of log-transformed $OD_{600}$ values in overlapping 5 h windows. In a chemostat, the slope $k$ of $\ln(OD_{600})$ over time corresponds to net growth, such that the specific growth rate was obtained as $\mu = D + k$, with $D = 0.15\ \mathrm{h}^{-1}$. For Ct, a replicate-specific estimate of $r_{Ct,A}$ was taken from the first sliding window, which captured the initial growth phase before substantial substrate depletion. For Oa, the earliest OD measurements contained an outlier that led to an overestimation of the initial growth rate. Therefore, a replicate-specific estimate of $r_{Oa,A,T}$ was obtained as the mean of the window-based growth rates over a broader time interval rather than from the single maximum estimate. The final values of $r_{Ct,A}$ and $r_{Oa,A,T}$ were then calculated as the averages across replicates. The parameter $r_{Ct,C}$, describing Ct growth on metabolite $C$, was set to 0.4 h$^{-1}$.

#### Substrate affinity constants

The Monod constants for acetate, $K_{Ct,A}$ and $K_{Oa,A}$, were inferred from monoculture chemostat $OD_{600}$ trajectories by least-squares fitting of the corresponding chemostat model (Fig. S1B–D). For each replicate, the growth-rate parameter $r$ was fixed to the value estimated as described above, and the yield $Y$ was approximated from the total increase in OD during growth divided by the initial acetate concentration of 7.5 mM. The initial biomass $N_0$ was set to the mean of the earliest OD measurements of each replicate, and the initial acetate concentration was set to $M_A = 7.5$ mM. The model was then simulated and $K$ was fitted by minimizing the difference between simulated and observed $OD_{600}$ trajectories. For Ct, fitting was performed over the interval from 1 to 12 h, whereas for Oa fitting was performed from 1 to 60 h. Replicate-specific estimates were obtained separately, and the final values of $K_{Ct,A}$ and $K_{Oa,A}$ were calculated as the averages across replicates.

To visualize the fitted difference in acetate affinity, we also plotted net growth rate $(d \ln N/dt)$ against $OD_{600}$ (Fig. S1E). Because OD increases as acetate is depleted, these plots provide a qualitative view of how quickly growth declines with resource consumption. Oa maintained positive net growth to higher OD values than Ct, consistent with its lower fitted acetate Monod constant.

The thiamine Monod constant $K_{Oa,T}$ was estimated from batch growth data of Oa grown across a thiamine gradient (SFig. S1F). To constrain this parameter, we used the 1 nM thiamine condition and simulated the corresponding batch dynamics with dilution set to zero ($D = 0$). All other parameters were fixed, and $K_{Oa,T}$ was fitted by least squares to the observed Oa $OD_{600}$ trajectory. The parameter $K_{Ct,C}$, describing Ct growth on metabolite $C$, was chosen manually for illustration.

For the thiamine production coefficient $q_{Ct,T}$, we performed a parameter sweep and selected a value that yields $\sim 100$ nM thiamine when Ct is grown alone in batch simulations (Fig. 1C). This choice is consistent with the observation that Oa reaches its full yield in Ct spent medium (Fig. 1A), and with the thiamine gradient experiment showing that $\geq 100$ nM thiamine restores full yield (Fig. S1F). For metabolite $C$, we chose $q_{Oa,C}$ such that Oa produces a total of 200 $\mu$M $C$ when grown alone in batch simulations (Fig. S5).
